# Single-Cell Metabolic Imaging Reveals Glycogen Driven-Adaptations in Endothelial Cells

**DOI:** 10.1101/2025.05.11.653328

**Authors:** Rahuljeet S. Chadha, Benjamin Yang, Dongqiang Yuan, Philip A. Kocheril, Shivansh Mahajan, Adrian Colazo, Naseeb K. Malhi, Joseph A. Ambarian, Xuejing Liu, Zhen B. Chen, Lu Wei

## Abstract

Endothelial dysfunction (ED) is a defining feature of diabetes mellitus (DM) and a key contributor to many metabolic and cardiovascular diseases. Endothelial cells (ECs) are known to be highly glycolytic and primarily rely on glucose to meet their energy demands. However, the role of glycogen metabolism in ECs remains poorly characterized due to a lack of suitable tools. Here, we utilize stimulated Raman scattering (SRS) microscopy to investigate subcellular glycogen metabolism in live ECs under stress conditions associated with highly prevalent diabetes and diabetic complications. We demonstrate that ECs exposed to a diabetes-mimicking milieu– high glucose and tumor necrosis factor (TNF-α)– divert excess glucose toward subcellular glycogen storage, and that this storage capacity is significantly enhanced by the inhibition of glycogen synthase kinase 3 (GSK3). Pulse-chase experiments uncover glycogen dynamics and reveal that glycogen is rapidly mobilized under glucose starvation, highlighting its role as an immediate energy reserve in ECs. We further extend the capabilities of SRS metabolic imaging to visualize glutamine and lactate metabolism for the first time, directly showcasing the reliance of ECs on these alternative carbon substrates during glucose deprivation. Our results indicate that ECs containing glycogen exhibit a reduced immediate metabolic demand for these gluconeogenic substrates in the absence of extracellular glucose. These findings suggest that glycogen may play a broader role beyond energy reserves in ECs by modulating stress-responsive metabolic adaptations and may offer potential therapeutic opportunities to address diabetes-induced ED and related cardiometabolic diseases.

## Introduction

Diabetes mellitus (DM) affects over 500 million people globally, with cardiovascular complications being a leading cause of disability and morbidity.^[1]^ A central driver of these complexities is endotheliopathy, or the dysfunction of endothelial cells (ECs), signified by a pro-inflammatory activation and a decreased nitric oxide production.^[2]^ ECs are strategically located, lining up the interior walls of all vascular networks. This positioning not only allows them to serve as a physical barrier between blood and tissues, but also as regulators of essential processes such as vascular homeostasis, transport of macromolecules, and recruitment of leukocytes.^[3],[4]^ When exposed to pathophysiological stimuli such as pro-inflammatory cytokines, hyperglycemia, and oxidative stress, ECs can become dysfunctional and contribute to a myriad of cardiometabolic diseases, including but not limited to atherosclerosis, pulmonary arterial hypertension (PAH), and diabetes.^[5–8]^ So far, the dysregulation of metabolites that drive ED progression remains largely unexplored at the subcellular level.^[9,10]^ This knowledge gap highlights the need to investigate the metabolic basis of ED under pathophysiological conditions such as diabetes, which can identify potential therapeutic strategies for ameliorating ED and related metabolic and cardiovascular diseases (CVDs).

The highly glycolytic nature of ECs makes them reliant primarily on glucose metabolism to meet their biosynthetic needs and energy demands.^[11]^ However, excess glucose and elevated inflammation, as in the case of diabetes, can induce endotheliopathy and exacerbate metabolic dysfunction.^[12]^ One potential metabolic outcome of excessive glucose is the formation of glycogen, a hyperbranched polymer of glucose.^[13]^ In cancer cells, high glucose has been observed to promote subcellular accumulation of glycogen, which in turn accelerates liver tumor initiation.^[14],[15]^ However, in ECs, the role of glycogen remains poorly understood.^[16],[17]^ Existing techniques face significant limitations for specific subcellular detection of glycogen: magnetic resonance imaging (MRI) with ^13^C-glucose lacks subcellular resolution^[18]^; positron emission tomography (PET) with [^18^F]-2-fluoro-2-deoxy-D-glucose (FDG) and fluorescence microscopy using 2-NBDG employ glucose tracers that are neither specific for glycogen^[13]^ nor capable of incorporation to glycolysis or glycogenesis pathways^[19,20]^; mass-spectrometry-based imaging methods are inherently destructive and incompatible with live-cell studies or longitudinal analyses.^[21]^

To address these challenges, stimulated Raman scattering (SRS) microscopy combined with deuterated metabolites offers a powerful, non-invasive method for specifically probing cellular metabolism in living systems by targeting the C–D vibration peaks corresponding to the metabolized biomass structures.^[22–25]^ In contrast to spontaneous Raman microscopy, SRS offers significantly enhanced sensitivity and rapid acquisition times that allow efficient mapping of these dynamic metabolites in living systems.^[26,27]^ Unlike conventional fluorescence microscopy, SRS imaging circumvents the need for bulky fluorophores or genetically encoded reporters, which can perturb biological systems and are prone to photobleaching.^[28]^ Compared to a purely label-free approach, the use of vibrational tags provides dynamic information on the incorporation, growth, and turnover of metabolites.^[21]^ Additionally, the compatibility of SRS with tissue imaging and linear concentration dependence for quantitative analysis render it a uniquely powerful tool for visualizing and studying subcellular metabolism with high spatial resolution in live cells, both in health and disease.

In this work, we leveraged SRS microscopy with deuterated glucose tracing to visualize glycogen metabolism in ECs. In a model of diabetic milieu (high glucose and TNF-a; HT),^[29]^ we captured the subcellular formation of glycogen pools in live ECs. HT induced significant glycogen accumulation, which was further amplified by inhibiting glycogen synthase kinase 3 (GSK3)– resulting in glycogen covering up to more than 50% of the cell area and highlighting the unexpectedly high glycogenic capacity of ECs under metabolic stress. Pulse-chase experiments showed dynamic glycogen turnover and spatial growth, supporting its role as a back-up energy reserve. To evaluate the functional significance of glycogen stores, glucose-starvation assays directly visualized the metabolic adaptation of ECs to lactate (Lac) and glutamine (Gln)– two alternative carbon substrates that fuel EC metabolism via truncated gluconeogenesis (GNG; reverse glycolysis) in the absence of extracellular glucose.^[30]^ This is also the first time that SRS metabolic imaging of lactate and glutamine was achieved. Notably, ECs with elevated glycogen levels showed reduced reliance on Lac and Gln metabolism, suggesting a role for glycogen in buffering the immediate metabolic demand in ECs during glucose-deprived conditions.

## Results

### Visualization of glycogen in live endothelial cells

We first established an *in vitro* model of EC dysfunction by exposing human umbilical vein endothelial cells (HUVECs) to high glucose and TNF-α (25 mM D-glucose and 5ng/mL TNF-α; termed “HT”) for three days to mimic hyperglycemia and chronic inflammation. As a normoglycemic and osmolarity control, cells were maintained in 25 mM D-mannitol (osmotic control; termed “NM”). As demonstrated previously,^[12,29]^ this combined HT treatment not only causes a strong induction of pro-inflammatory response but also an endothelial-mesenchymal transition phenotype, evidenced by the marker gene expression and cell morphology (**Figure 1a**). Concomitant with the induction of pro-inflammatory genes *ICAM1* and *VCAM1*, quantitative PCR (qPCR) revealed an increased expression of *GYS1*, an isoform of glycogen synthase (GS)^[31]^ and the key rate-limiting enzyme of glycogen synthesis,^[32]^ in HT-exposed ECs (**Figure 1b**). This led us to examine glycogenesis in dysfunctional ECs using live-cell SRS imaging. Specifically, we utilized a glucose isotopologue (d_7_-glucose) as a functional substitute for glucose, where all carbon atoms are labeled by deuterium. Unlike FDG and 2-NBDG, d_7_-glucose is fully metabolized through native glucose metabolism pathways, allowing it to label the major biomolecules such as lipids, proteins, nucleic acids, and glycogen, for chemical mapping (**Figure 1c**). Each of these biomacromolecules is characterized by a distinct C–D Raman spectral signature arising from their local chemical environments.^[13],[33]^

**Figure 1.**
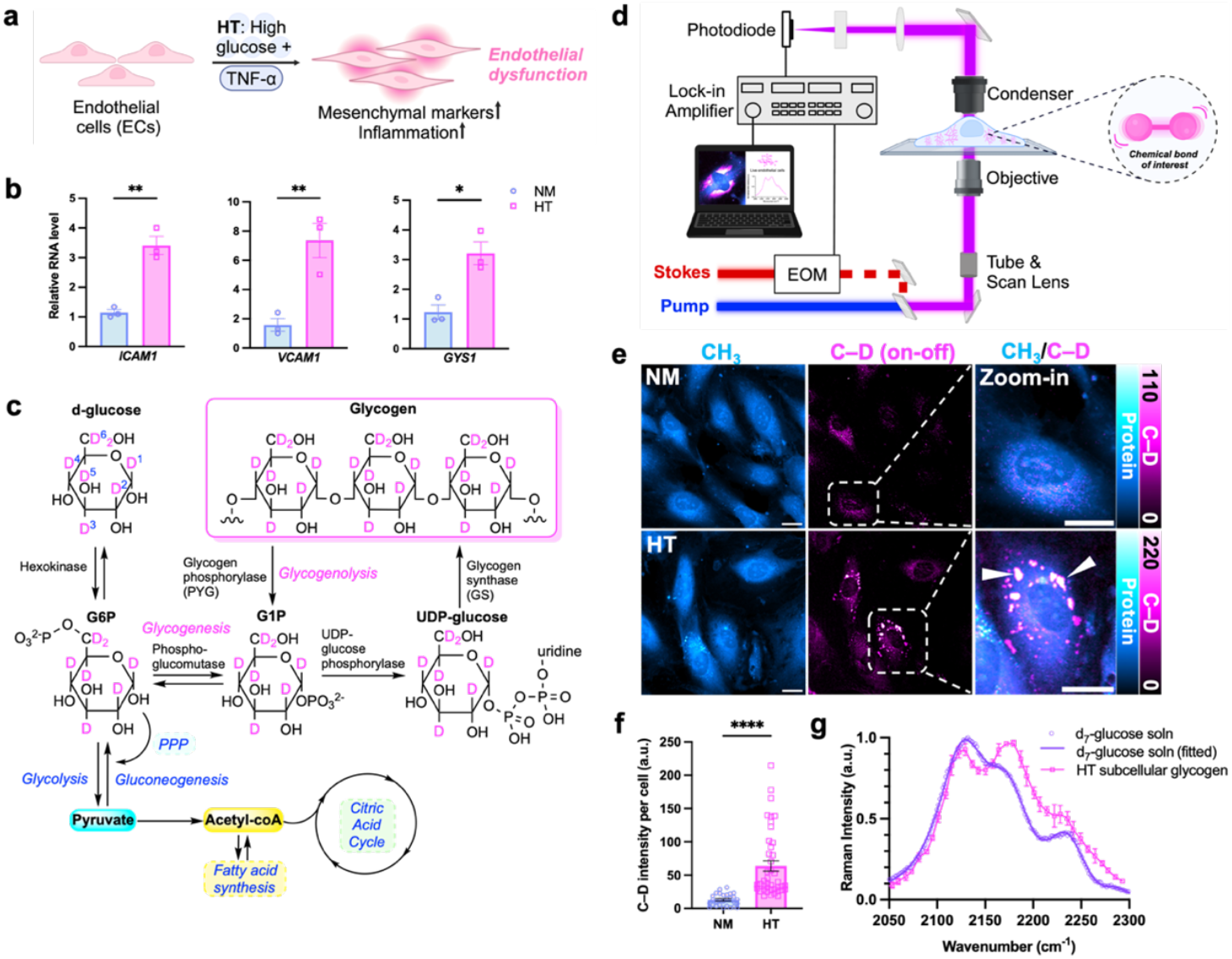
Visualization of subcellular glycogen in endothelial cells under HT. **(a)** Illustration of ECs exposed to high glucose and TNF-α (HT) and undergoing dysfunctional changes, characterized by an increase in inflammation and mesenchymal markers; **(b)** mRNA levels of *ICAM1, VCAM1* and *GYS1* in (osmotic control, NM) vs HT-treated HUVECs for 3 days; **(c)** Scheme of metabolic incorporation pathways of d_7_-glucose, including the glycogen synthesis (glycogenesis) pathway. PPP= Pentose Phosphate Pathway; **(d)** Schematic of the SRS microscopy configuration. EOM: Electro-optic modulator; **(e)** Representative SRS images of live HUVECs treated with NM or HT containing d_7_-glucose for 3 days, targeted at the –CH_3_ (protein) and C–D (on-off resonance) channel. The third column represents merged (CH_3_/C–D) channels for the zoomed-in view of ECs (boxed) in the corresponding C–D images. White arrowheads represent subcellular glycogen pools. Scale bar: 20 µm. **(f)** Quantitative analysis of SRS C-D signal per cell for NM and HT-treated cells (n=29 cells for NM and 42 cells for HT). **(g)** Normalized spontaneous Raman spectrum of d_7_-glucose solution (soln) (purple circles, 300 mM; purple solid line, fitted data). Normalized SRS spectrum of subcellular glycogen from HT-treated ECs (magenta squares with solid line). Data in all bar plots (**b, f**) is presented as mean ± SEM from at least three independent experiments. Statistical significance was analyzed using two-tailed unpaired Student’s t-tests. *p<0.05, **p<0.01, ****p<0.0001.

SRS imaging (**Figure 1d**) clearly revealed the appearance of subcellular glycogen pools labeled with d_7_-glucose as localized puncta (**Figure 1e**, white arrowhead) in HT-exposed ECs, indicated by a pronounced increase in the overall C–D signal per cell (**Figure 1f**). Because glycogen is composed of polymerized glucose units, its identity can be directly confirmed in live cells using hyperspectral SRS (hSRS). The spectral features of the observed subcellular puncta match those of d_7_-glucose, with all major peaks (e.g. 2120 and 2172 cm^-1^) consistent with known glucose solution spectra (**Figure 1g**).^[13],[33]^

Since downstream glucose-derived biomolecules like lipids and proteins produce overlapping but distinct C–D peaks, we applied two complementary chemometric methods– spectral phasor^[34]^ and least absolute shrinkage and selection operator (LASSO)^[35]^– to specifically resolve and unmix C–D signal arising from glycogen in dysfunctional ECs (**Figure S1**). These methods confirmed that the enriched spots in the C–D channel indeed correspond to glycogen. Moreover, these two methods provide a robust approach for glycogen phenotyping in biosystems.

To further validate these findings, we performed correlative SRS imaging with an established glycogen histochemical staining method, Periodic Acid-Schiff (PAS) staining using bright-field microscopy. Our results demonstrated a high correlation of identified glycogen deposits between SRS imaging and PAS staining (**Figure S2**). Additionally, we verified that the observed glycogen reservoirs did not arise from the aggregation of d_7_-glucose by detecting glycogen in ECs exposed to HT containing unlabeled D-glucose using PAS staining (**Figure S3**). However, we caution that fixation, as required in PAS detection, induced extracellular vesicles (fi-EVs) enriched with glycogen (**Figure S3–S4**), representing a fixation artifact consistent with our previous observations in cancer cells.^[13]^ Such vesicles are absent in all live-cell measurements, underscoring the rigor and advantage of performing unbiased live-cell investigations. Collectively, we successfully visualized and characterized C–D imaging of glycogen in ECs. To the best of our knowledge, this is the first report of subcellular glycogen visualization in live ECs.

### Pharmacological modulation of GSK3 reveals a high capacity for glycogen storage in ECs

We next sought to modulate glycogen accumulation by targeting glycogen synthase kinase 3 (GSK3), a serine/threonine kinase that phosphorylates GS to inhibit glycogen synthesis^[36]^ (**Figure 2a**). In its active form, GSK3 phosphorylates glycogen synthase (GS) to make it less active for glycogen synthesis. Inhibition of GSK3 causes dephosphorylation of GS, thereby making it more active for glycogen synthesis. We treated HT-exposed ECs with two small-molecule inhibitors of GSK3, CHIR-99021^[37]^ or SB-216763^[38]^ (hereafter CHIR and SB, respectively). Compared to ECs exposed to HT alone, HT+CHIR resulted in a striking increase in d_7_-glucose-tracked glycogen accumulation, with glycogen pools covering over 50% of the cellular area (**Figure 2b-c**). Similar results were confirmed by using 6,6-d_2_-glucose (hereafter d_2_-glucose), another glucose isotopologue, bearing fewer deuterium labels (**Figure 2d-e**) and a different Raman spectral signature (**Figure S5**). Such drastic increase in glycogen storage was consistent with the effect of SB, the other GSK3 inhibitor (**Figure S6**).

**Figure 2.**
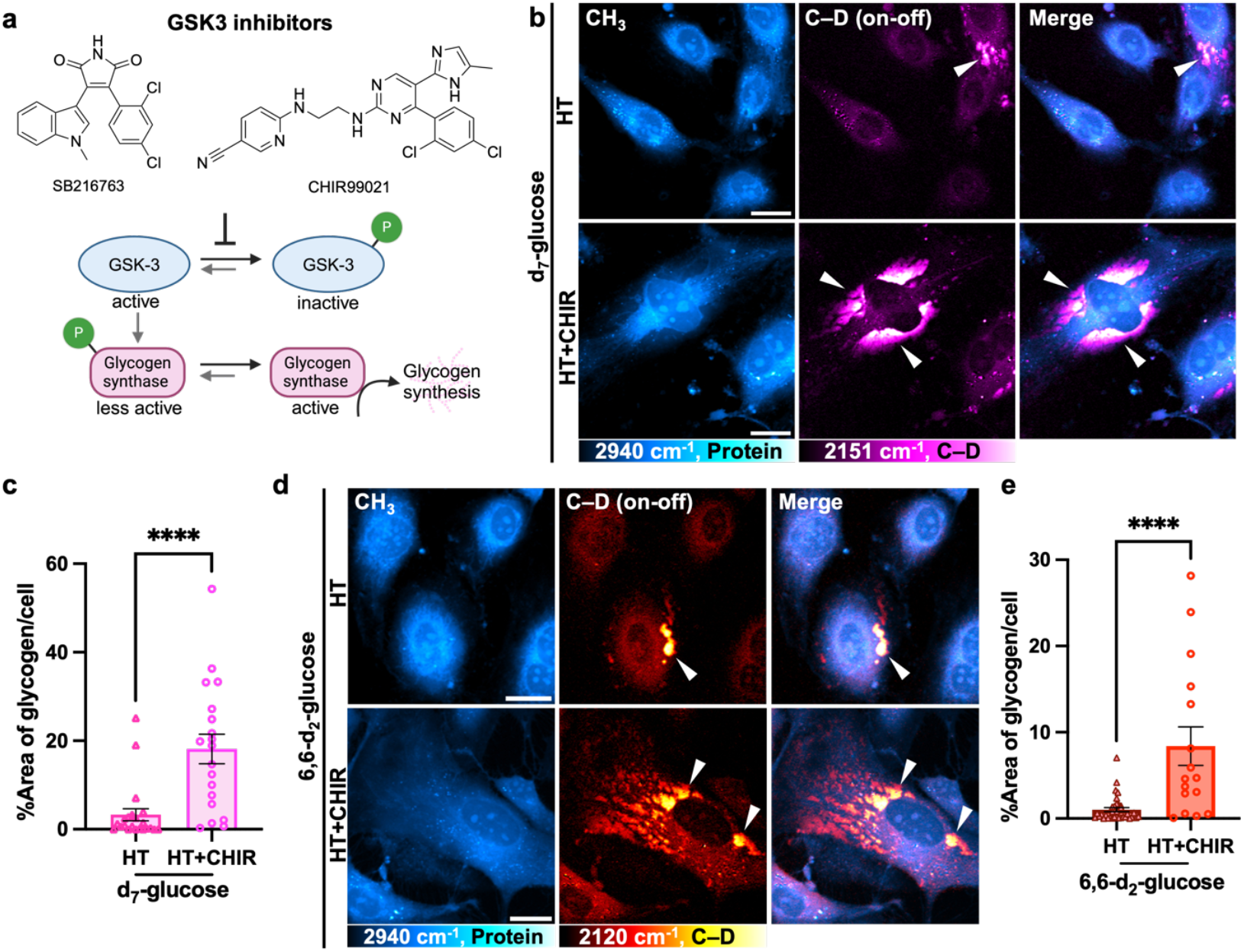
Pharmacological inhibition of GSK3 reveals a high capacity for glycogen storage in ECs. **(a)** Chemical structures of small molecule inhibitors of GSK3, SB216763 and CHIR99021; and the mechanistic scheme for inhibition of GSK-3 to activate glycogen synthesis. **(b)** Left to right: representative live-cell SRS images at the –CH_3_ (protein), C– D (on-off), merged (CH_3_/C–D) channels for HUVECs treated with HT, or HT+CHIR containing d_7_-glucose for three days; **(c)** Quantification of the percentage of glycogen area per cell from HT and HT+CHIR-treated ECs (n=22 and 19 cells, respectively.); **(d)** Left to right: representative live-cell SRS images at the –CH_3_ (protein), C–D (on-off), merged (CH_3_/C–D) channels for HUVECs treated with HT, or HT+CHIR containing 6,6-d_2_-glucose for three days. White arrowheads represent subcellular glycogen pools; **(e)** Quantification of the percentage of glycogen area per cell from HT and HT+CHIR-treated ECs (n=36 and 16 cells, respectively). Data (**c, e**) presented as mean ± SEM from at least three independent experiments. Statistical significance was analyzed using two-tailed unpaired Student’s t-tests. ****p<0.0001. Scale bar: 20 µm.

In parallel to single-cell imaging, we performed a bulk glycogen assay using enzyme-coupled luminescence reactions, which confirmed a ∼2.3-fold total increase in glycogen concentration in ECs treated with HT+CHIR compared to HT alone (**Figure S7, Table S1**). Our results imply a surprisingly high storage capability for ECs under GSK3 inhibition. By leveraging GSK3 inhibition to enhance glycogen accumulation, this model also provides a robust platform to investigate the spatial growth and subcellular distribution of glycogen clusters in greater detail.

### Dynamics of glycogen formation, spatial organization, and degradation in endothelial cells

To advance the understanding of glycogen synthesis (glycogenesis) and degradation (glycogenolysis) in ECs, we performed longitudinal SRS imaging in live ECs pulsed with HT and d_7_-glucose. We observed increased glucose incorporation into biomass in ECs over time, including progressive accumulation of subcellular glycogen reservoirs (**HT; Figure 3a, S8**). While a previous study based on stable isotope labeling and mass spectrometry reported mobilization of glycogen reserves in bulk and lysed ECs under hypoglycemia,^[39]^ our imaging approach directly visualized glycogen breakdown at the subcellular level in live ECs: our chase experiment, following a glucose starvation chase, observed rapid glycogenolysis, with most of the subcellular pools depleted by 48 hours in ECs pre-exposed to HT (**HT; Figure 3b, S8**).

**Figure 3.**
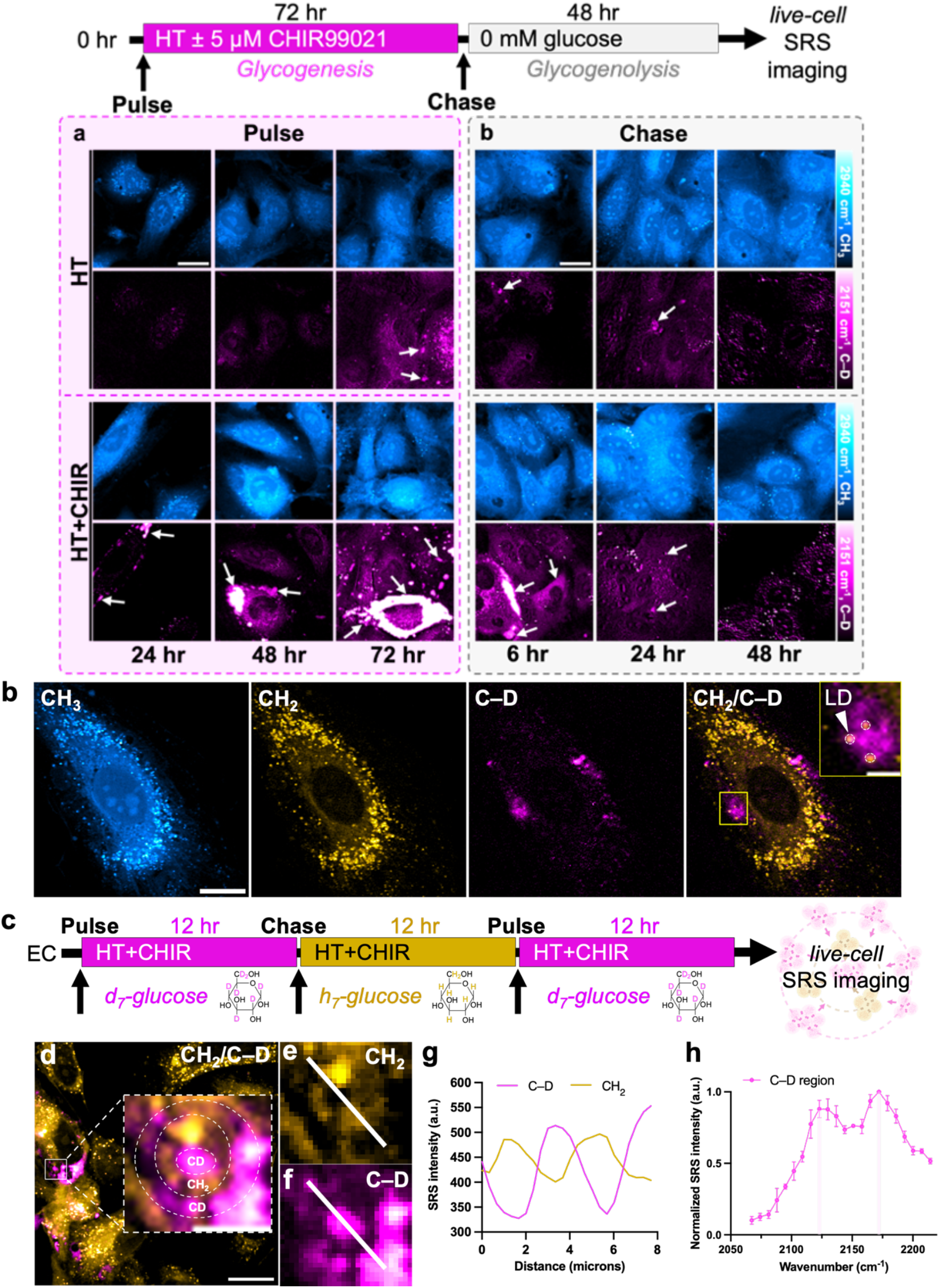
Glycogen dynamics, spatial organization, and degradation in endothelial cells. **(a)** Experimental design for the pulse-chase experiment to probe glycogenesis and glycogenolysis. Representative SRS images targeting the –CH_3_ (protein) and C–D (on-off) channel after 24-, 48- and 72-hr of pulse with HT or HT+CHIR-treated HUVECs (top) and after 6-, 24- and 48-hr of chase in glucose-free media (bottom). White arrows indicate glycogen pools; **(b)** Multi-channel SRS images at the –CH_3_ (protein), –CH_2_ (lipid) and C– D (on-off) channels for HUVECs treated with HT+CHIR for 8-hr. Scale bar: 20 µm. Overlay images of CH_2_ and C–D with an inset indicate lipid droplets (LDs; yellow circles) colocalized with glycogen clusters. Inset scale bar: 1 µm; **(c)** Design of two-color glycogen experiment; **(d)** Overlay of CH_2_ and C–D channel. Scale bar: 20 µm. Inset: a zoomed-in region of a bullseye pattern of glycogen formed from alternating concentric rings (dotted white circles) of glucose isotopologues. Inset scale bar: 5 µm; **(e-f)** CH_2_ (e) and C–D (f) image of the inset in (d), indicating a glycogen ring formed from unlabeled H-glucose (e) and the two concentric rings of glycogen formed from d_7_-glucose (f); **(g)** Intensity profile across the white solid line in (e) and (f); **(h)** Normalized SRS spectrum across the white-line in (f) confirming glycogen composition. Data are plotted as mean ± SEM.

We further investigated glycogen turnover under HT+CHIR conditions, which markedly increased glycogen storage capacity (**HT+CHIR; Figure 3a, S8**). Similar to HT, glycogen pools in HT+CHIR also underwent rapid glycogenolysis under glucose deprivation with both conditions depleting their glycogen pool by 48 hours (**HT+CHIR; Figure 3b, S8**). Taken together, these results indicate the highly dynamic nature of these glycogen reservoirs and support their role as immediate energy reservoirs in ECs under glucose-limited conditions.

Building on these findings, we next investigated glycogen initiation and localization by performing multi-channel SRS imaging in HUVECs treated with HT+CHIR for 8 hours. Interestingly, we observed glycogen cluster initiation at multiple sites in close proximity to lipid droplets (LDs) (**Figure 3b**). In parallel experiments, we noted a striking increase in the LD abundance in ECs treated with HT compared to NM, consistent with other models of EC dysfunction.^[40–42]^ In other closely related cell types, such as brown hepatocytes, glycogen dynamics have been linked to driving lipid droplet (LD) biogenesis during cellular differentiation.^[43]^ While further investigation is needed, these observations raise the intriguing possibility of potential metabolic crosstalk between LDs and glycogen dynamics in dysfunctional ECs.

To visualize glycogen cluster assembly and expansion in real space and time, we performed a two-color pulse-chase experiment using sequential labeling of d_7_-glucose and normal glucose media. HUVECs were first pulsed with d_7_-glucose for 12 hours, chased with unlabeled D-glucose for another 12 hours, and then pulsed again with d_7_-glucose for 12 hours during CHIR treatment (**Figure 3c**). SRS microscopy revealed a bullseye pattern of concentric rings, reflecting alternating incorporation of labeled and unlabeled glucose within glycogen clusters. This pattern suggests that glycogen protrusions expand radially from a central origin and can span over several microns under hyperglycemic stress (**Figure 3d-h**). While diffraction-limited SRS microscopy (∼400 nm spatial resolution)^[21]^ does not resolve ultrastructural features such as β- or α-particles seen by transmission electron microscopy (TEM),^[44]^ it uniquely captures the spatial dynamics of glycogen synthesis and organization in live ECs.

### SRS metabolic imaging of glutamine and lactate reveals metabolic shifts in ECs containing glycogen

The glycogen accumulation in ECs under HT conditions led us to hypothesize that this adaptation may confer a survival advantage during glucose deprivation. Surprisingly, HUVEC viability remained largely unaffected even in glycogen-deficient HUVECs (NM) after 72 hours of glucose starvation (**Figure S9**). In contrast, cervical cancer-derived HeLa cells which are also highly glycolytic,^[45]^ showed significantly decreased viability under the same conditions (**Figure S9**). Building on our observation that glycogen serves as an energy reserve in ECs and exhibits rapid turnover (*vide supra*), we hypothesized the elevated presence of glycogen may initially affect the metabolic demand for alternative carbon sources when extracellular glucose is scarce. To investigate the metabolic adaptations that enable ECs to sustain their energy requirements in glucose-deprived environments, we focused on glutamine and lactate metabolism, two key metabolites known to undergo truncated GNG in ECs to generate lower glycolytic intermediates that support serine and glycerophospholipid biosynthesis pathways.^[30]^

Glutamine, the most abundant freely circulating amino acid in the body, can fuel metabolism in various cell types including cancer cells under glucose stress.^[46]^ It contributes to the tricarboxylic (TCA) cycle, nucleotide and nonessential amino acid (NEAA) synthesis,^[47]^ and can fuel GNG by converting to glutamate and subsequently α-ketoglutarate (**Figure 4a**).^[48]^ In ECs, glutamine is also essential for vascular expansion and proliferation.^[49,50]^ To visualize glutamine metabolism, we coupled SRS microscopy with d_5_-glutamine, which exhibits a distinct C–D Raman peak at 2165 cm^-1^ (**Figure 4b, Figure S10a**). HUVECs were pulsed for three days with NM (negligible glycogen), HT (moderate glycogen), or HT+CHIR (high glycogen), followed by a 24-hour chase with 5 mM d_5_-glutamine in glucose-free medium.

**Figure 4.**
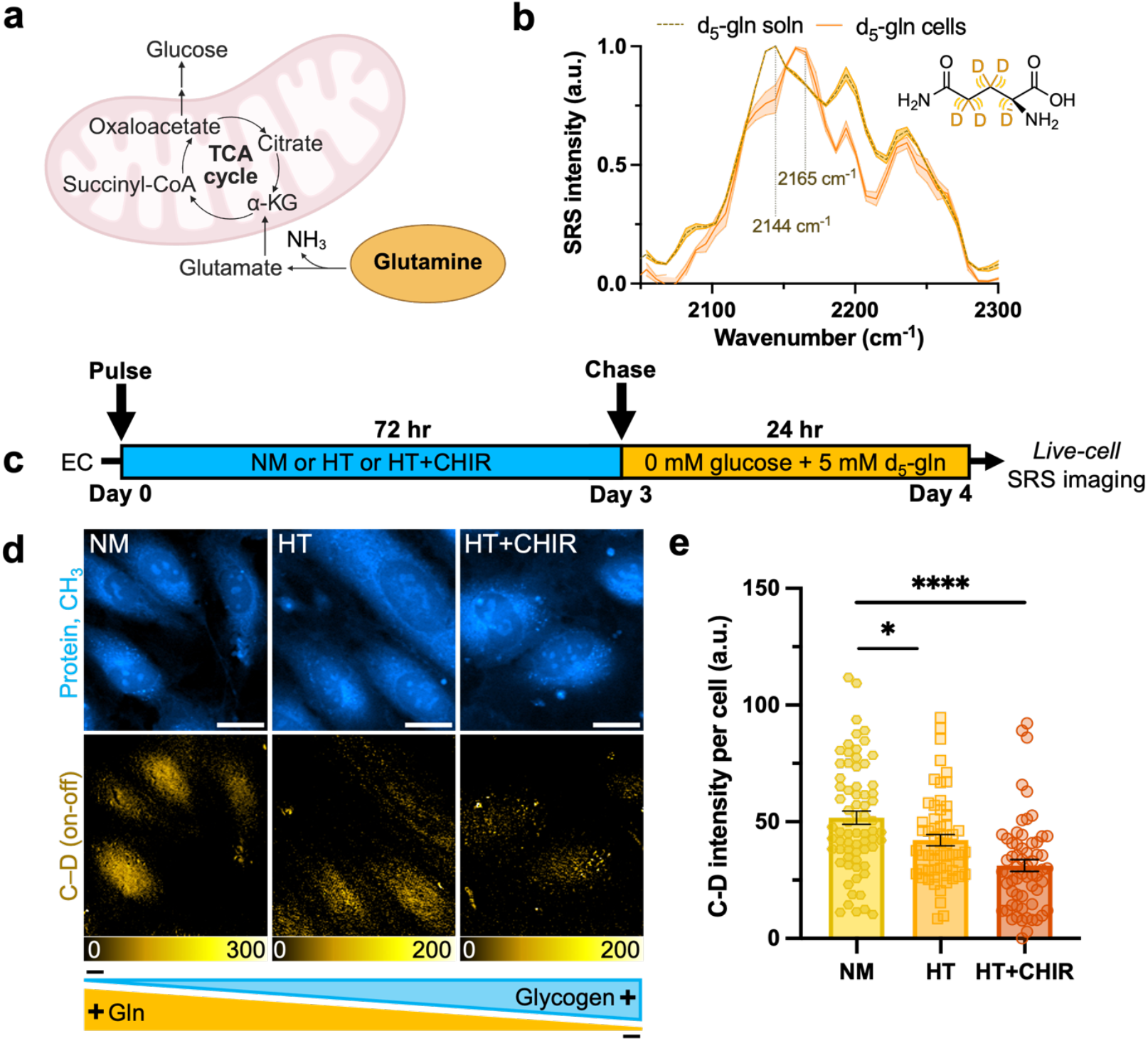
SRS microscopy reveals a decreased metabolic reliance on glutamine in ECs containing glycogen. **(a)** Schematic illustration of the gluconeogenic pathway of glutamine; **(b)** Normalized SRS spectra of 100 mM d_5_-gln solution, and of HUVECs incubated with 30 mM d_5_-gln for three days; **(c)** Schematic of pulse-chase experiment to probe glutamine metabolism; **(d)** Representative SRS images at the –CH_3_ and C–D (on-off) for NM, HT and HT+CHIR. Scale bar: 20 µm; **(e)** Quantitative analysis of SRS signal of C–D bonds per cell (n=69, 59, 62 cells for NM, HT, and HT+CHIR, respectively). Data are presented as mean ± SEM from at least three independent experiments. Statistical significance was analyzed using two-tailed unpaired Student’s t-tests. ****p<0.0001,*p<0.05.

Strikingly, in the absence of extracellular glucose, ECs containing moderate to high contents of glycogen (both HT and HT+CHIR) showed reduced glutamine incorporation, compared to NM-treated cells, indicating a lower metabolic demand for glutamine in the presence of glycogen (**Figure 4c-d**). As a control, we tested glutamine metabolism in all three conditions after glycogen stores were mostly exhausted by performing a longer chase in glucose-free medium and found that this difference diminished over time (**Figure S11**). This result aligns with the metabolic shift seen in the liver where GNG increases to maintain euglycemia (normal glucose levels) as glycogen is exhausted.^[51],[52]^

To determine whether this decrease in glutamine signal was due to altered protein synthesis, we traced d_5_-phenylalanine metabolism, an essential amino acid, as a proxy. SRS imaging targeted at the C–D channel at 2298 cm^-1^ (**Figure S12a**) showed no significant differences in phenylalanine-derived protein synthesis among NM, HT, and HT+CHIR (**Figure S12b-c**). These results support the previous hypothesis that the reduction in glutamine signal in glycogen-rich conditions should be mostly contributed from glutamine metabolism itself, and not due to a reduced demand for protein synthesis.

We next moved to examine lactate metabolism– another key alternative carbon source.^[53]^ Once considered a metabolic waste product in mammals, lactate is now recognized as a major circulating metabolic fuel, especially during glucose deprivation.^[54]^ As a critical GNG precursor via pyruvate (**Figure 5a**),^[55],[56]^ lactate is traditionally studied using genetically encoded fluorescent sensors^[57,58]^ which face limitations in their dynamic range, genetic accessibility, and applicability in primary cells like ECs. To address these challenges, we employed SRS microscopy coupled with d_3_-lactate, targeting the C–D channel at 2129 cm^-1^ (**Figure 5b, Figure S10b**). HUVECs pretreated with NM, HT, or HT+CHIR for 3 days were chased in 30 mM d_3_-lactate for 24 hours in glucose-free media. Similar to glutamine, lactate incorporation was reduced in glycogen-rich ECs (HT and HT+CHIR) compared to NM (**Figure 5c-d**). This difference was also mitigated over time as glycogen was consumed (**Figure S13**), further reinforcing the idea that glycogen modulates the immediate metabolic adaptations of ECs to alternative fuels during glucose starvation. Notably, this also marks the first time that glutamine and lactate metabolism have been visualized in live cells using SRS imaging.

**Figure 5.**
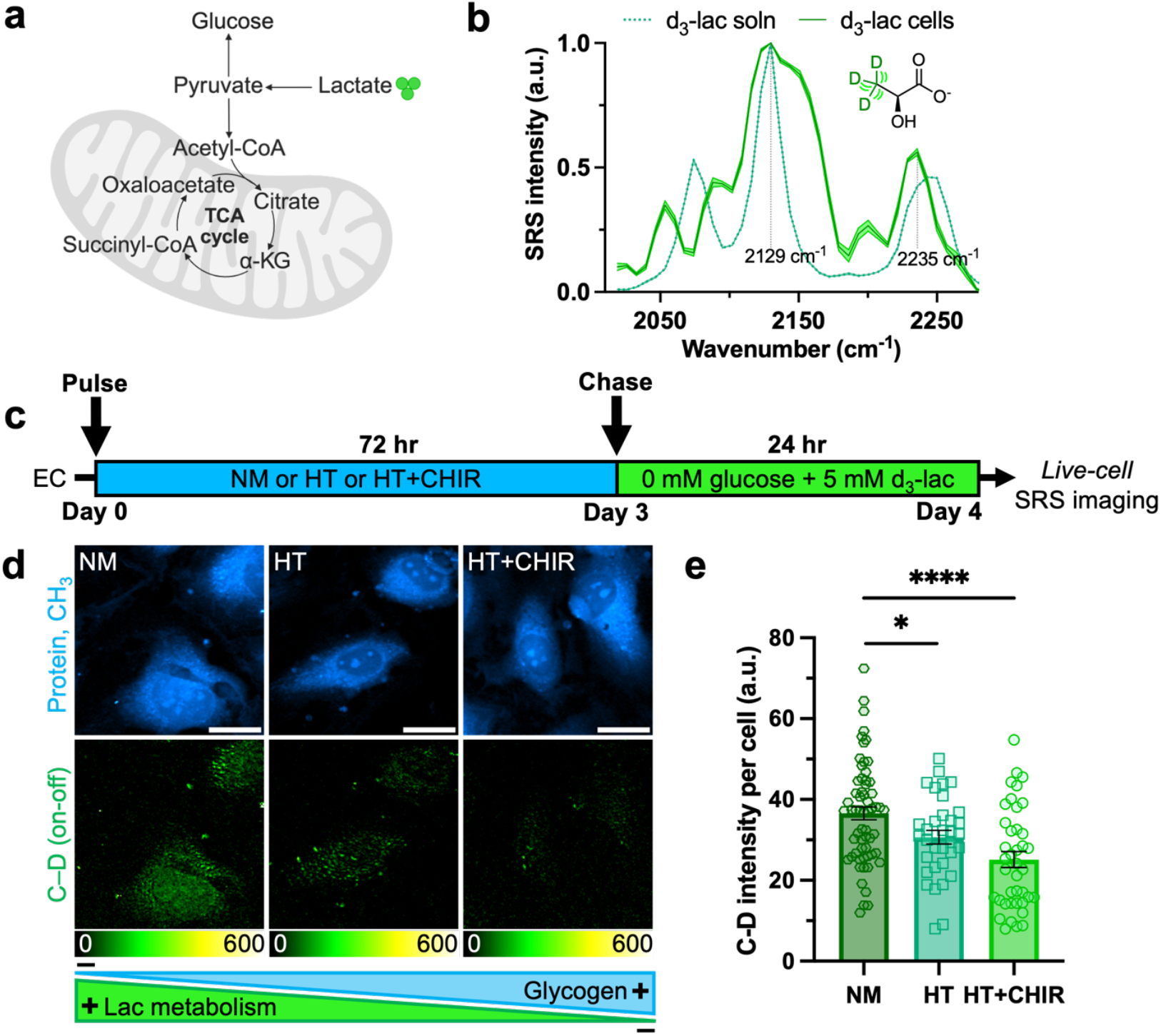
SRS microscopy reveals a decreased metabolic reliance on lactate in ECs containing glycogen. **(a)** Schematic illustration of the gluconeogenic pathway of lactate in cells; **(b)** Normalized SRS spectra of 30 mM d_3_-lac solution, and of HUVECs incubated with 30 mM d_3_-lac for three days; **(c)** Schematic of pulse-chase experiment to probe lac metabolism; **(d)** Representative SRS images at the –CH_3_ and C–D (on-off) for NM, HT and HT+CHIR. Scale bar: 20 µm; **(e)** Quantitative analysis of SRS signal of C–D bonds per cell (n=62, 34, 40 cells for NM, HT and HT+CHIR, respectively). Data are presented as mean ± SEM from at least three independent experiments. Statistical significance was analyzed using two-tailed unpaired Student’s t-tests. ****p<0.0001,*p<0.05.

## Discussion

Diabetes-induced vascular dysfunction is driven by metabolic stressors such as hyperglycemia and hypoxia, which disrupt regulation of metabolites such as glucose across various cell types.^[59],[60]^ However, the role of glycogen metabolism in endothelial dysfunction remains poorly understood. Our study addresses this gap by revealing the dynamic nature of glycogen metabolism in ECs under diabetes-related metabolic stress. By leveraging longitudinal SRS microscopy with non-perturbative deuterium labeling, we provide direct visualization of subcellular glycogen pools in live ECs and track their rapid turnover in response to hyperglycemia and glucose deprivation. Furthermore, SRS imaging of glutamine and lactate demonstrates that glycogen reserves initially modulate ECs’ reliance on these alternative carbon sources under glucose starvation. These findings offer new insights into the metabolic adaptations that accompany diabetes-associated endothelial dysfunction.

Future work includes exploring the generality of these findings in other EC types, such as microvascular (MVECs) and sinusoidal ECs (SECs), as well as primary ECs harvested from clinically relevant tissue samples. Given the ability of ECs to crosstalk with other cell types including immune cells,^[61]^ perivascular cells,^[62]^ and adipocytes,^[63]^ it would be compelling to explore whether EC-derived glycogen can be transferred to other cells to better understand metabolic crosstalk. In addition to in vitro systems, non-invasive visualization of glycogen and associated metabolic adaptations in vivo remains a key goal. Beyond glutamine and lactate, future studies should also explore the role of other key GNG substrates such as pyruvate, glycerol, alanine, etc.^[64]^ in driving metabolic adaptations in ECs.

While our study provides new insights, certain limitations should be addressed in future investigations. For instance, although lactate supplementation was necessary to probe its metabolism, supraphysiological concentrations could also influence endothelial cell biology, potentially triggering endothelial-to-mesenchymal transition (EndoMT) under metabolic stress.^[65]^ Additionally, the current sensitivity limits of SRS imaging constrain the ability to resolve the complete metabolic fate of vibrational probes. To gain a more comprehensive understanding of substrate-specific utilization and fates, complementary approaches such as mass spectrometry-based metabolic flux analysis (MFA)^[66]^ with higher detection sensitivities may be necessary.

Overall, our work shows that dysfunctional ECs respond to hyperglycemic conditions by forming glycogen pools that serve as short-term energy backup during metabolic stress. Beyond this conventional role, these dynamic stores also initially influence the metabolic reliance on alternate carbon sources during the absence of extracellular glucose. Moreover, our demonstration of SRS metabolic imaging of lactate and glutamine in living cells unlocks new possibilities for visualizing the underlying metabolic changes of diabetes-induced endothelial dysfunction and related pathologies.

## Materials and Methods

### Cell Culture, Metabolic Labeling and Imaging

HUVECs (passages 5-8; Cell Applications, Cat. #200-05n) were tested negative for mycoplasma contamination and cultured at 37°C with 5% CO_2_ in commercial endothelial cell growth media (ECGM; Cell Applications, Cat. #211-500), supplemented with 10% FBS (Corning, Cat. #35-015-CV) and 100× antibiotic-antimycotic (Gibco, Cat. #15240062). HeLa cells (ATCC, CCL-2) were cultured in DMEM (Corning, Cat. #10-017-CV) with the same supplementation as above. For SRS imaging experiments, HUVECs were seeded on glass coverslips pre-coated with attachment factor solution (Cell Applications, Cat. #123-500) for 2 hours. The HT condition was induced by adjusting the final concentration to 25 mM D-glucose (Thermo Fisher, Cat. #A16828-36), d_7_-glucose (Cambridge Isotope Laboratories, Cat. #DLM-2062), or 6,6-d_2_-glucose (Cambridge Isotope Laboratories, Cat. #DLM-349), along with 5 ng/mL TNF-α (Gibco, Cat. #PHC3015). The NM condition included addition of 25 mM mannitol (Thermo Fisher, Cat. #125340025) as an osmolarity control in ECGM containing 5 mM baseline D-glucose. For GSK3 inhibition, 5 µM CHIR-99021 (MedChemExpress, Cat. #HY-10182), or 5 µM SB-216763 (MedChemExpress, Cat. #HY-12012) was added concomitantly with the HT condition containing labeled or unlabeled glucose for the specified duration.

For the glycogen dynamics experiments, HUVECs were treated with HT (using DMSO as a vehicle) or HT+CHIR in medium containing labeled d_7_-glucose for the specified duration. To probe glycogenolysis, cells were subsequently chased in custom glucose-free ECGM (Cell Applications, Cat. #211AG-500) for the respective time points before live-cell SRS imaging.

For the two-color glycogen experiment, HUVECs were first pulsed with HT (containing labeled d_7_-glucose)+CHIR for 12 hours, then chased with HT (containing unlabeled glucose)+CHIR for 12 hours, and finally pulsed again with HT (containing labeled d_7_-glucose)+CHIR for another 12 hours before SRS imaging.

For the glutamine and lactate chase experiments, HUVECs were treated with NM, HT, or HT+CHIR containing unlabeled glucose for three days, followed by a chase with the labeled probes added at the desired concentration for 24 hours in custom media containing 0 mM glucose, 0 mM unlabeled L-glutamine and 0 mM unlabeled lactate (Cell Applications, Cat. #211AG-500). The labeled probes used for metabolic imaging were 5 mM 2,3,3,4,4-d_5_-L-glutamine (Cambridge Isotope Laboratories, Cat. #DLM-1826) and 30 mM 3,3,3-d_3_ sodium L-lactate (Cambridge Isotope Laboratories, Cat. #DLM-9071) for glutamine and lactate, respectively. The metabolic label used for phenylalanine imaging was d_5_-L-phenylalanine (Cambridge Isotope Laboratories, Cat. #DLM-1258).

### SRS Microscopy

The SRS microscopy system was configured as described previously.^[67]^ **Figure 1d** depicts the corresponding microscope configuration. Briefly, images were acquired with an 80 µs pixel dwell time, achieving an image acquisition speed of 8.52 s per frame for a 320x320-pixel field of view at a 0.497 µm/pixel resolution. The pump beam wavelength was set to 791.3 nm for the protein (–CH_3_ channel, 2940 cm^-1^), 797.3 nm (–CH_2_ channel, 2845 cm^-1^), 844.0 nm (C–D channel for d_7_-glucose, 2150 cm^-1^), 846.2 nm (C–D channel for 6,6-d_2_-glucose, 2120 cm^-1^), 842.9 nm (C–D channel for d_5_-glutamine, 2167 cm^-1^), 845.6 nm (C–D channel for d_3_-lactate, 2129 cm^-1^), 833.7 nm (C–D channel for d_5_-phenylalanine, 2297 cm^-1^) and 867.0 nm (off-resonance, 1837 cm^-1^). For hSRS, the wavelength of the pump laser was tuned from 833.5 nm to 853.5 nm with a step size of 0.5 nm unless indicated otherwise. All images were analyzed and color-coded using ImageJ software. For C–D quantification, on-resonance images were processed by subtracting the corresponding off-resonance images to result in the C–D (on-off) images. Individual cells were then manually segmented, and the SRS intensity per cell was quantified following background subtraction. MS Excel, GraphPad Prism, and MATLAB (R2024b, MathWorks) were used for data processing and plotting.

### Spectral Phasor Analysis

The hSRS sweeps across the range 2067–2235 cm^-1^ (25 images with a step size of ∼7 cm^-1^) were imported into ImageJ as a stack. A maximum intensity projection was created and spectral phasor analysis was performed using a spectral phasor plugin (http://www.spechron.com/Spectral%20Phasor-Download.aspx) as described previously.^[34]^ Segmentation of the phasor plot was performed manually to discretely map glycogen.

### LASSO

The same set of aforementioned hSRS images were also unmixed using least absolute shrinkage and selection operator (LASSO)^[35],[68]^ in MATLAB (R2024b, MathWorks) on a per-pixel basis following nonuniform background removal and non-local means denoising in ImageJ.^[69,70]^

### Spontaneous Raman Microscopy

Spontaneous Raman spectra were acquired using a Horiba Xplora plus upright confocal Raman spectrometer with a 532 nm YAG laser at 12 mW through a 100x, 0.9 N.A. objective (MPLAN N, Olympus) with a 500 µm hole, and a 100 µm slit. Data were collected with an integration time of 10 s and averaged 10 times using LabSpec6 software. The background signal from the solvents was subtracted from the spectra of the target analytes followed by baseline correction, and normalization.

### Periodic Acid-Schiff (PAS) Staining

HUVECs were incubated under HT treatment (containing labeled or unlabeled glucose) for three days, fixed with 4% paraformaldehyde for 15 minutes, and washed three times using Dulbecco’s phosphate-buffered saline (DPBS, Gibco, Cat. #14190144). Cells were then treated with 1% periodic acid (Sigma Aldrich, Cat. #3951) for 30 mins at room temperature and washed five times using deionized water. After incubation with Schiff’s reagent (Sigma Aldrich, Cat. #3952016) for 30 mins and additional washes with deionized water, PAS-stained cells were imaged on an Olympus FV3000 microscope using a color CMOS camera (Thorlabs, CS165CU) in tandem with SRS microscopy.

### Cell Viability Assay

For the viability assay under glucose starvation, cells cultured on 24-well plates were switched to glucose-deficient media and chased for 1-3 days. Relative cell viability was calculated using trypan blue staining and normalized to the control that consisted of cells grown in the corresponding baseline glucose-containing media.

### RNA Extraction and Quantitative PCR

RNA extraction was performed using Trizol reagent (Ambion, Cat. #15596018). RNA concentration was measured using Nanodrop spectrophotometer (Thermo Fisher). PrimeScriptTM RT Master Mix contained both Oligo-dT and random hexamer primers. Quantitative PCR was performed using SYBR Green Master Mix (Bio-Rad) following the manufacturer’s suggested protocol, using the Bio-Rad CFX Optus PCR System (Bio-Rad). The primer sequences used are listed in **Table 1**. Relative expression levels were determined using the comparative CT method to normalize target genes to *18S* internal controls.

**Table 1.**
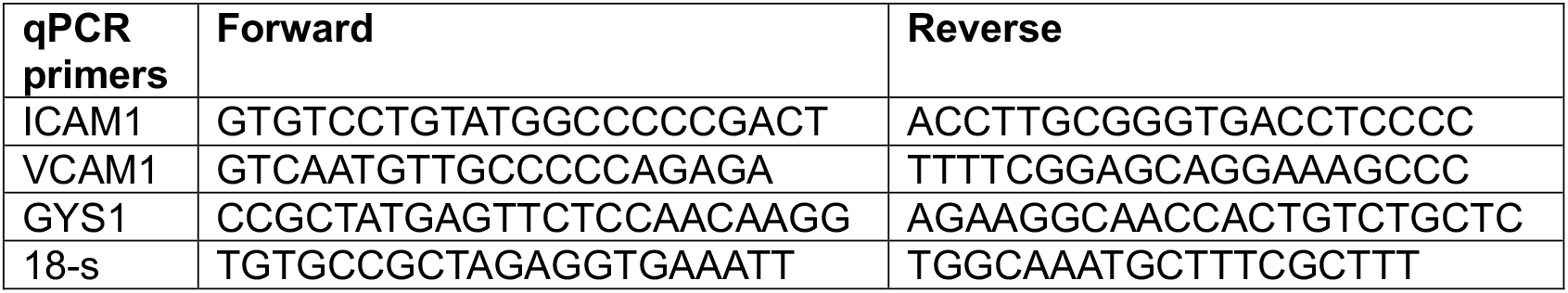
List of primer sequences used for qPCR.

### Glycogen Luminescence Assay

In preparation for glycogen detection, HUVECs cultured on 6-cm dishes were treated with HT, or HT+CHIR for three days. Glycogen detection was performed using the Glycogen-Glo™ Assay kit (Promega, Cat. #J5051) following manufacturer’s protocol. Luminescence was measured on a SpectraMax M3 multi-mode microplate reader (Molecular Devices) in relative light units (RLU). The sample buffer with no cells was used as a negative control. A control with no glucoamylase was also used to detect residual glucose in the samples. All samples were run as both technical and biological triplicates. Glycogen and glucose calibration curves were plotted in the concentration ranges of 0 to 20 µg/mL and 0 to 2.5 µg/mL, respectively. Glycogen and glucose concentrations in samples were calculated using the corresponding calibration plots in MS Excel and plotted in Prism GraphPad.

## Supporting Information

Supporting Information is available from the Wiley Online Library or from the author.

## Acknowledgements

We acknowledge fruitful discussions with Dr. Lei Jiang and Dr. Ryan Leighton. We thank the Newman Lab (Caltech) for the use of their microplate reader. Select figures were created using Biorender.com. R.S.C. acknowledges the support received from Pittcon/ACS Analytical Graduate Fellowship. P.A.K. is grateful for financial support from a National Science Foundation Graduate Research Fellowship (DGE-1745301) and a Hertz Fellowship. A.C. acknowledges support from NSF Graduate Research Fellowship (DGE-2139433). Z.B.C. is supported by R35 HL171550 from National Institute of Health. X.L. and N.K.M. are supported by postdoctoral fellowships from American Heart Association (24POST1195441 and 25POST1365287). L.W. is a Heritage Principal Investigator supported by the Heritage Medical Research Institute and also acknowledges support from a CZI dynamic imaging grant.

## Author Contributions

R.S.C., L.W., Z.B.C. conceptualized and designed the experiments. R.S.C., B.Y., D.Y., J.A.A. and S.M. conducted the experiments. N.K.M and X.L. assisted in conducting experiments. A.C. assisted in data acquisition with SRS imaging. P.A.K. assisted in data processing. The manuscript was written by R.S.C., L.W. and Z.B.C. with input from all authors.

## Data Availability Statement

The data that support the findings of this study are available from the corresponding author upon reasonable request.

