## Supporting information for "Single-Cell Metabolic Imaging Reveals Glycogen Driven-Adaptations in Endothelial Cells"

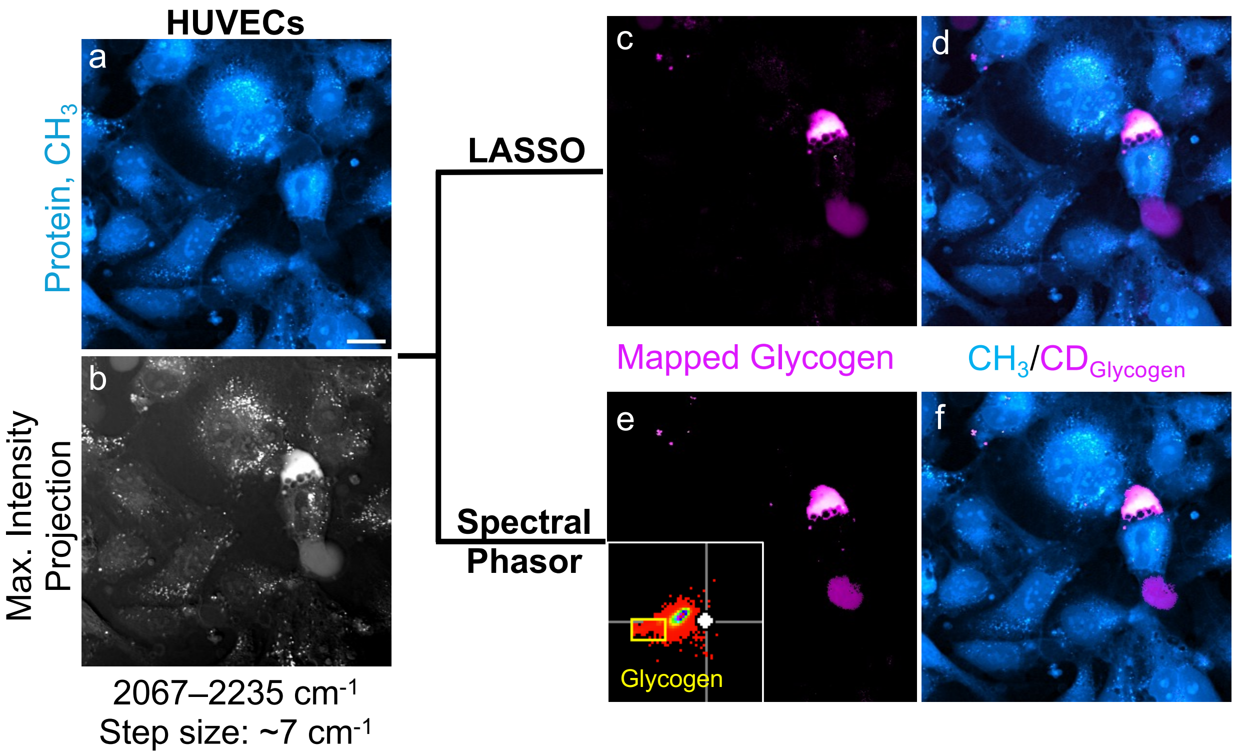


**Figure S1. Chemometric approaches for glycogen phenotyping in ECs. (a)** SRS image targeting the –$CH_3$ channel (protein) in ECs under hyperglycemic conditions; **(b)** Maximum intensity projection of a hSRS stack comprising 25 images acquired across 2067-2235 $cm^{-1}$ with a step size of 7 $cm^{-1}$ used for chemometric analysis; **(c)** Mapped glycogen image generated using LASSO ($CD_{Glycogen}$); **(d)** Overlay of $CH_3$, and $CD_{Glycogen}$ image derived from LASSO analysis; **(e)** Mapped glycogen image generated using spectral phasor analysis ($CD_{Glycogen}$). The inset shows the phasor plot where the boxed area (yellow) represents pixels attributed to glycogen; **(f)** Overlay of $CH_3$, and $CD_{Glycogen}$ image generated using spectral phasor analysis. Scale bar: 20 μm.

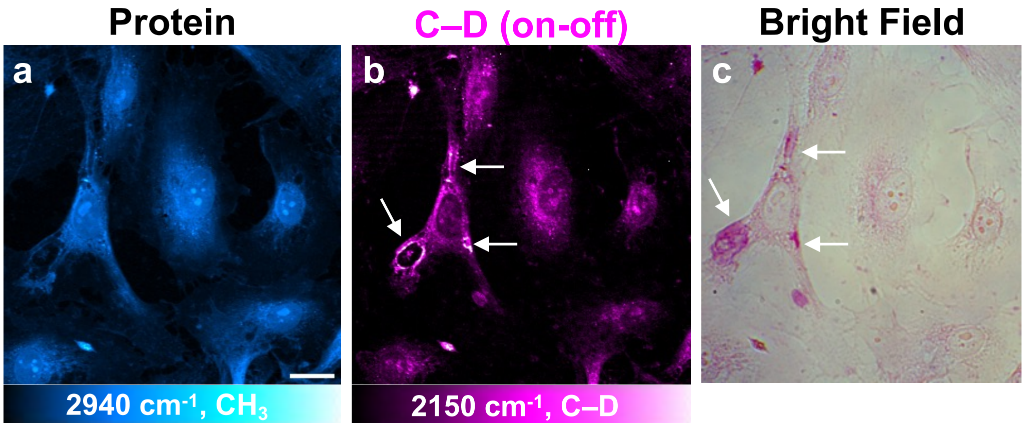


**Figure S2. Correlative PAS staining of glycogen pools in ECs. (a, b)** Representative SRS images targeted at the $–CH_3$ (protein) and C–D (on–off) channels, respectively, for HT-treated HUVECs; **(c)** Bright field image of the same set of cells after PAS staining. White arrows represent glycogen pools. Scale bar: 20 μm.

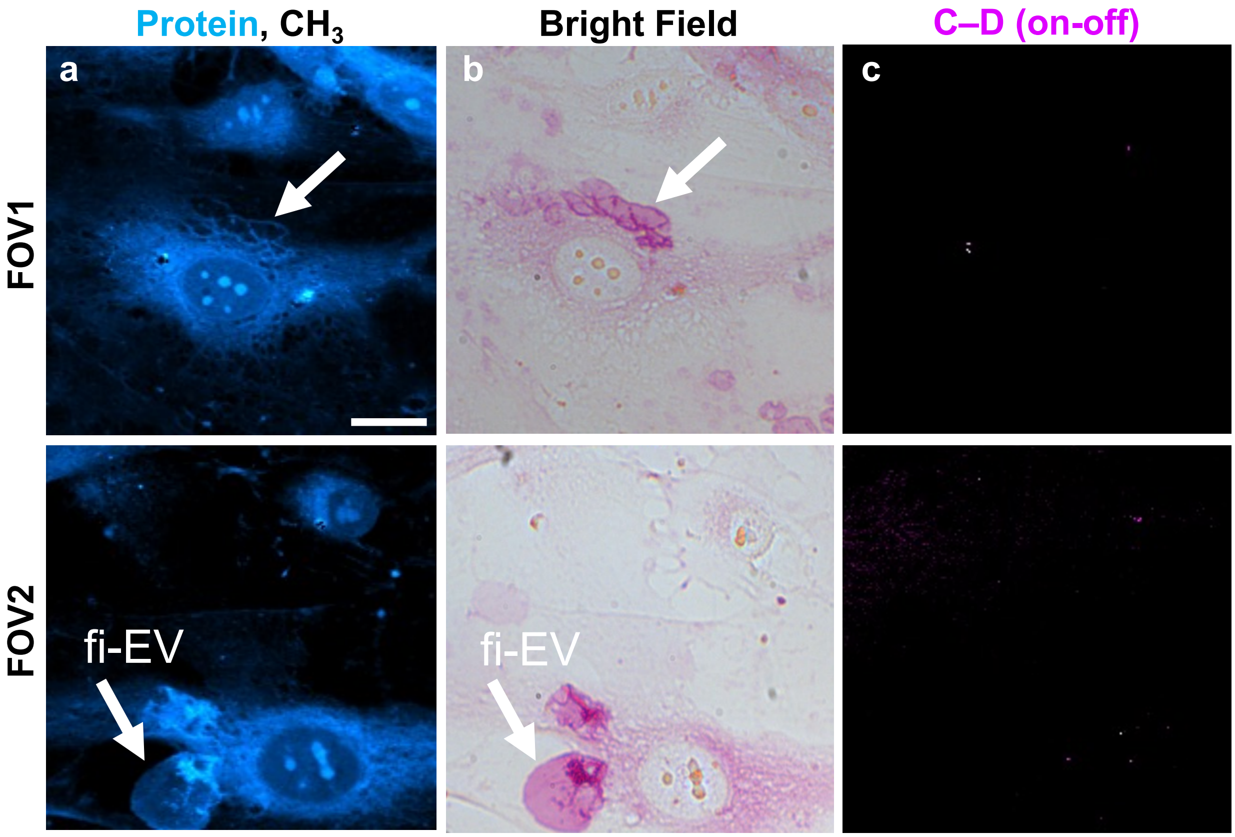


**Figure S3. Glycogen pools in ECs are not an artifact of glucose isotopologues.** **(a)** Representative SRS image targeted at the $–CH_3$ (protein) for HT-treated HUVECs containing unlabeled glucose; **(b)** Bright field image of the same set of cells after PAS staining; **(c)** C–D images (on–off) of the same corresponding field of views (FOVs). White arrows represent glycogen pools; fi-EV = fixation-induced extracellular vesicle. Scale bar: 20 μm.

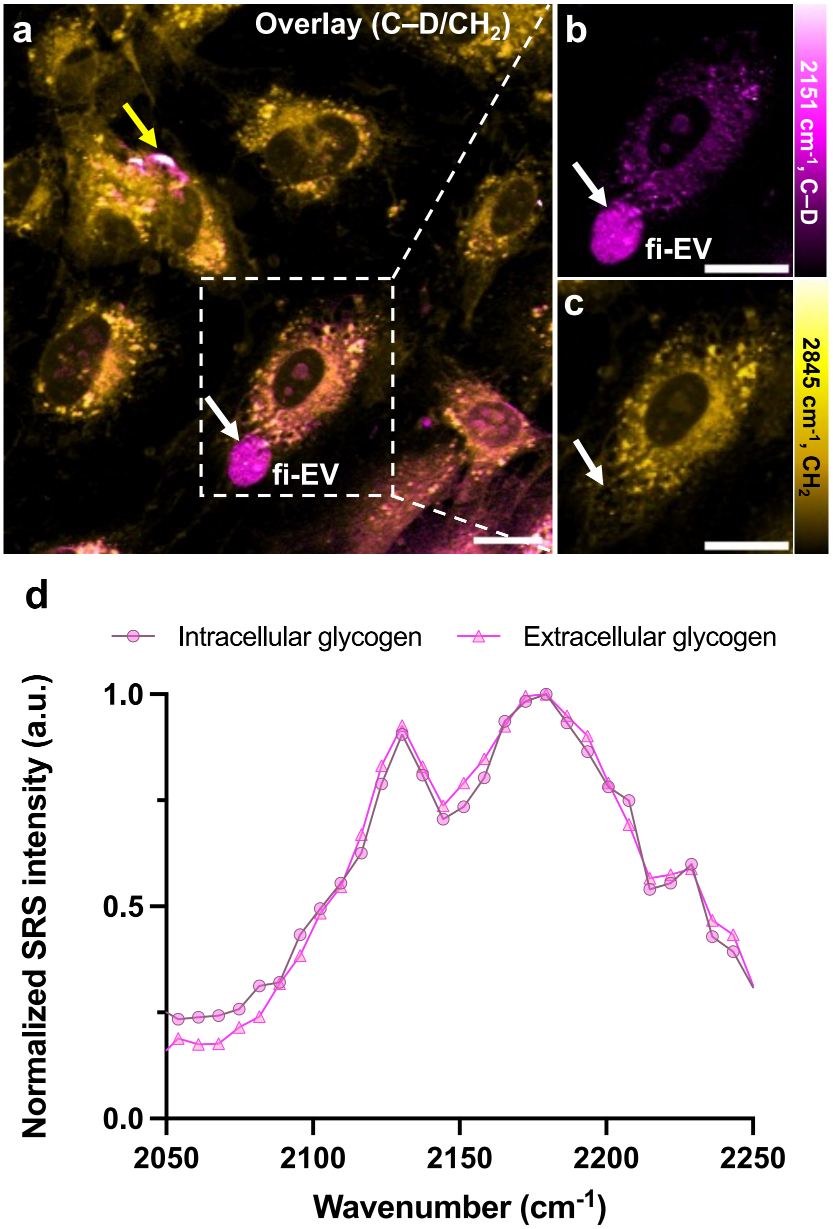


**Figure S4. PFA fixation induces glycogen-enriched extracellular vesicles. (a)** Representative overlay image (C–D/$CH_2$ channels) of fixed HUVECs treated with HT using $d_7$-glucose for three days. Yellow arrow represents intracellular glycogen reserves, while the white arrow indicates the extracellular glycogen in fixation-induced EVs (fi-EVs) after PFA fixation; **(b)** Zoomed-in image of the boxed region in (a), targeted at the C–D channel (on–off) indicating glycogen-enriched fi-EV (white arrow); **(c)** Corresponding zoomed-in $CH_2$ (lipid) image of the boxed region in (a); **(d)** Normalized SRS spectrum of intracellular and extracellular glycogen regions from (a). Scale bar: 20 μm.

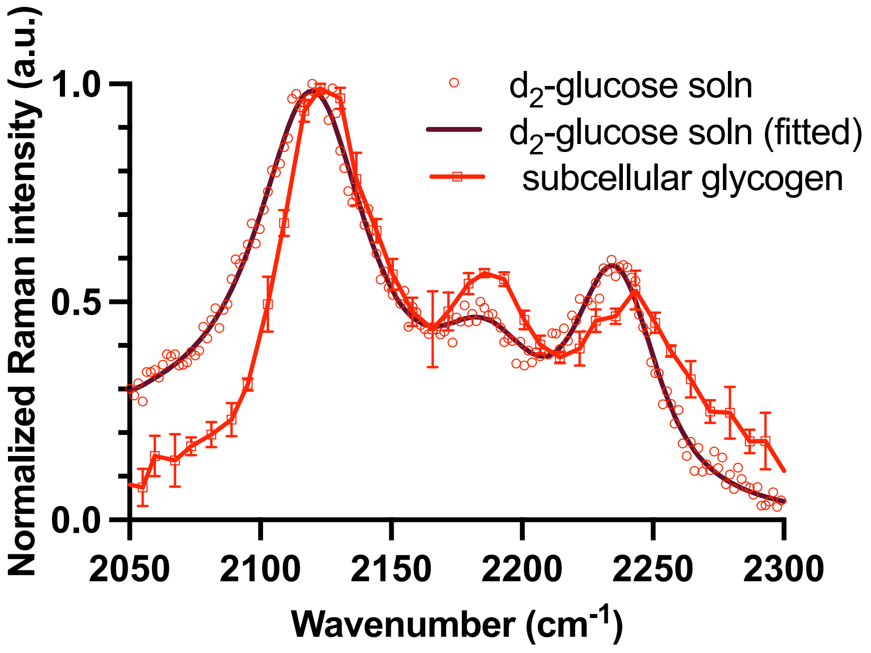


**Figure S5. Raman spectra of glucose isotopologue, 6,6-d$_2$-glucose.** Normalized spontaneous Raman spectrum of 100 mM solution (soln) of 6,6-d$_2$-glucose in water (red circles) with corresponding fitted data (maroon solid line). Normalized SRS spectrum of subcellular glycogen from HT-treated ECs incubated with d$_2$-glucose (red squares with solid line). Data are presented as mean ± SEM.

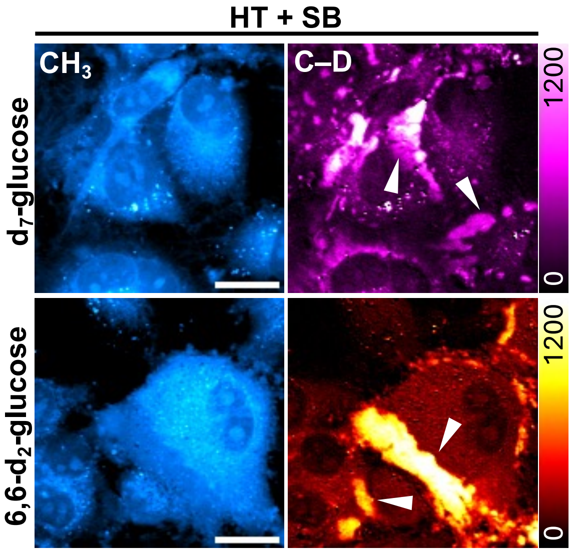


**Figure S6. GSK3 inhibition using an alternative small molecule inhibitor, SB-216763.** Representative SRS images targeting the $–CH_3$ (protein) and C–D (on-off) channel for live HUVECs treated with HT+SB incubated with either $d_7$-glucose, or $d_2$-glucose. White arrowheads represent subcellular glycogen pools. Scale bar: 20 μm.

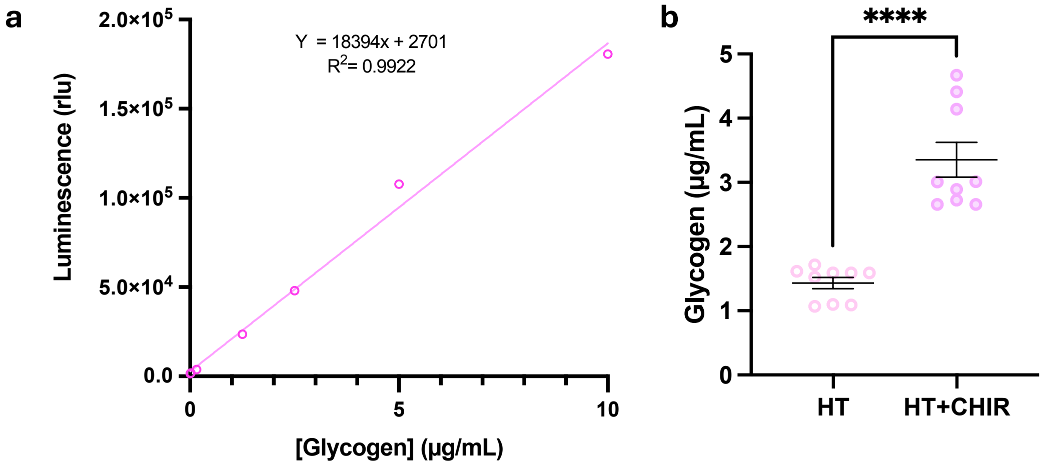


**Figure S7. Glycogen detection in bulk ECs using luminescence. (a)** Calibration curve for glycogen with different concentrations (μg/mL) measured by luminescence (rlu); **(b)** Glycogen concentration (μg/mL) in ECs treated with HT, or HT+CHIR for three days, quantified using the calibration curve in (a) (n=9 technical replicates from three independent experiments). Statistical significance was analyzed using two-tailed unpaired Student's t-test, ****$p<0.0001$. Data are presented as mean ± SEM.

**Table S1. Glycogen concentration detected in bulk HUVECs using luminescence.** Mean concentration of glycogen (μg/mL) for the corresponding treatments detected using the luminescence assay (in Figure S7) and their standard error of mean (SEM). The analytical sensitivity ($\gamma$) of the assay is calculated to be 18,394 rlu/(μg/mL).

| Condition | Glycogen conc. (μg/mL) | SEM (μg/mL) |
|---|---|---|
| HT | 1.43 | 0.08 |
| HT+CHIR | 3.35 | 0.27 |

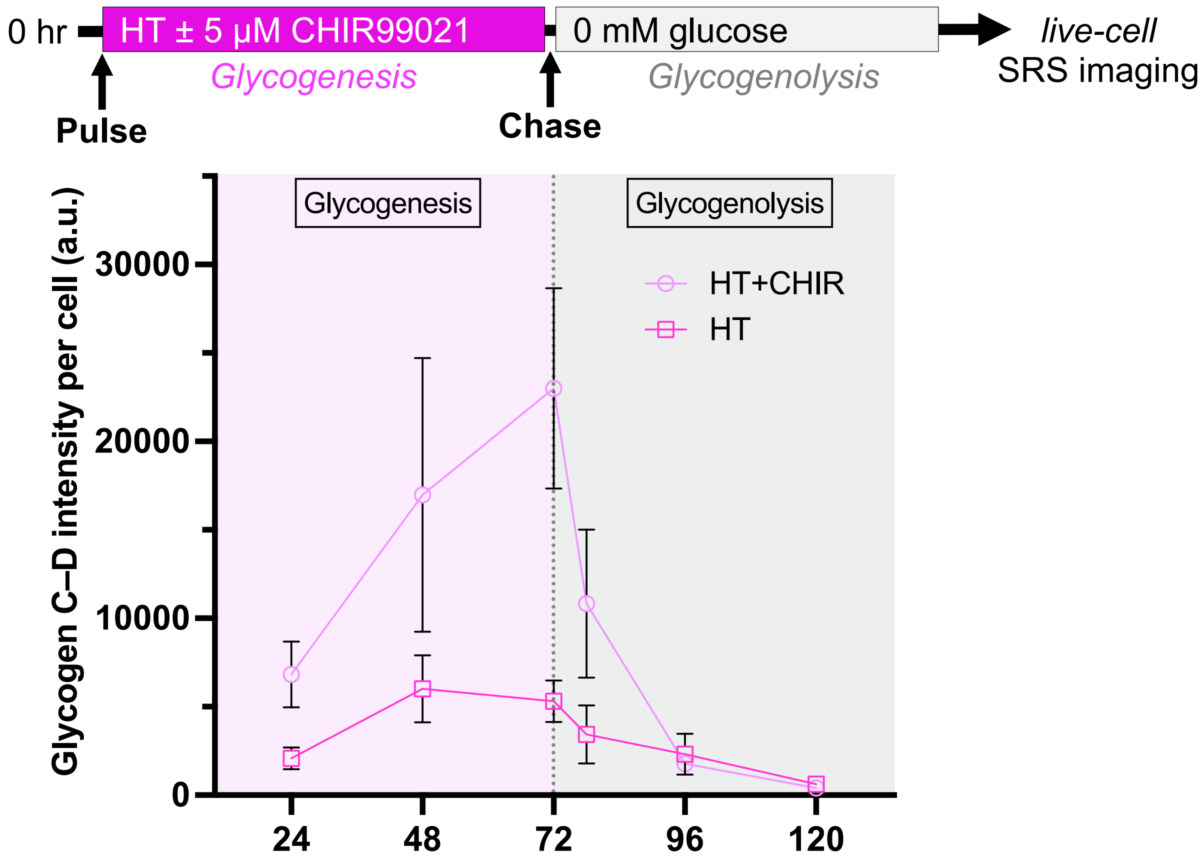


**Figure S8. Longitudinal imaging of glycogen metabolism in ECs.** SRS intensities of segmented glycogen C–D (on-off) per cell pulsed with HT, or HT+CHIR for 72 hr, and chased in glucose-free media for 48 hr. Data are presented as mean ± SEM from at least three independent experiments.

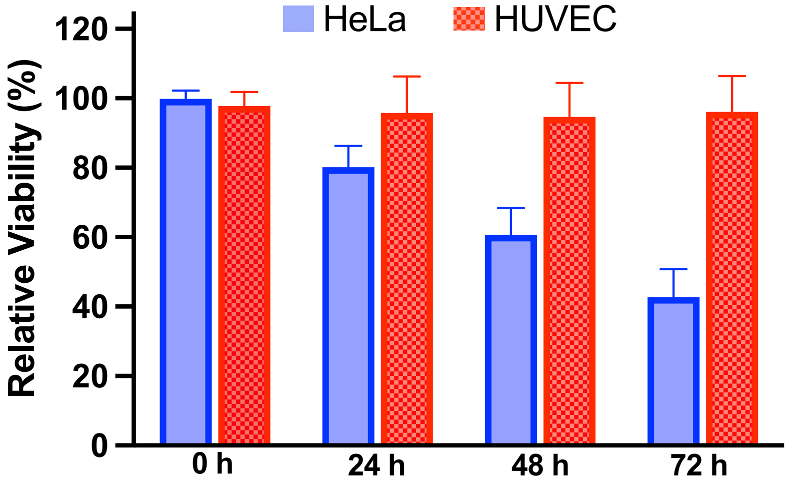


**Figure S9. Viability assay for HeLa and HUVEC under glucose starvation.** Relative viability of HeLa and HUVECs cultured in glucose-deficient media for 72 hours. Data are presented as mean ± SEM.

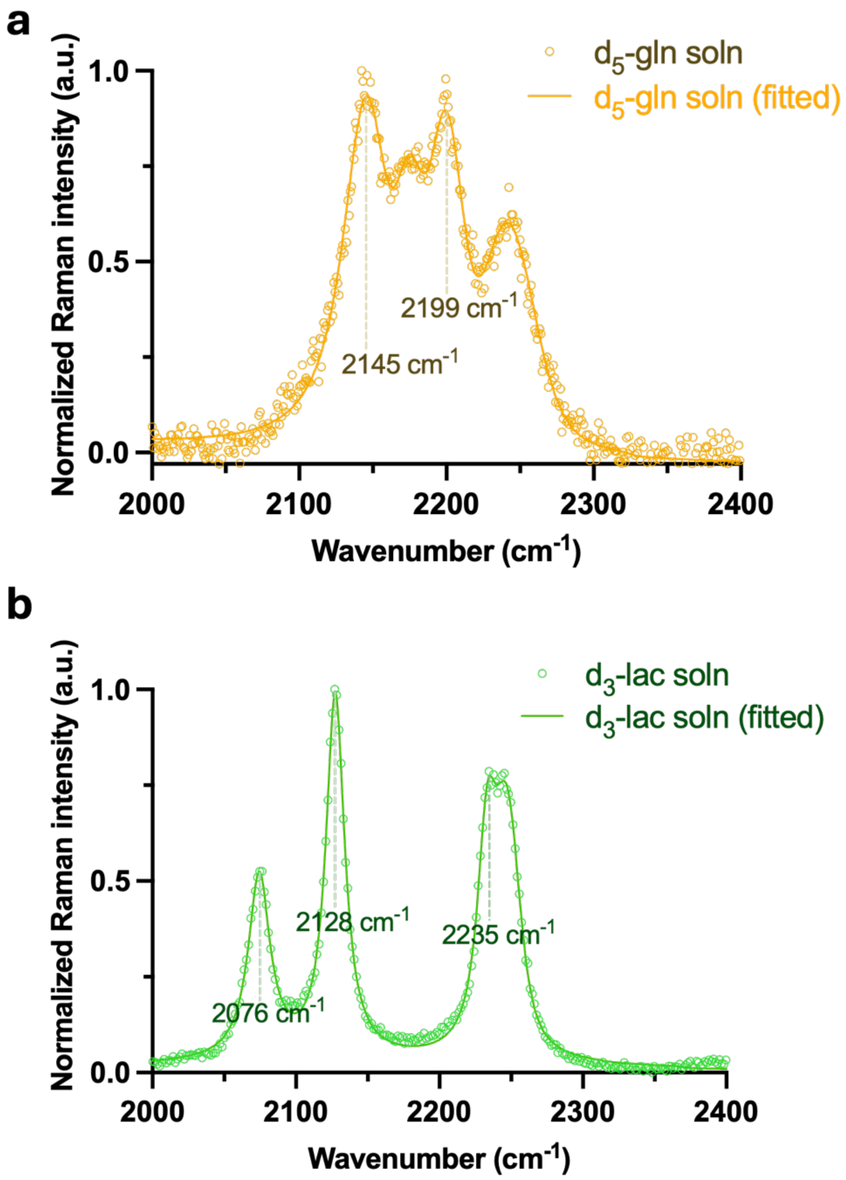


**Figure S10**. Spontaneous Raman spectra of **(a)** 200 mM of $d_5$-glutamine solution (soln) in water (experimental, yellow circles; fitted, yellow solid line) and **(b)** 25 mM $d_3$-lactate solution (soln) in water (experimental, green circles; fitted, green solid line).

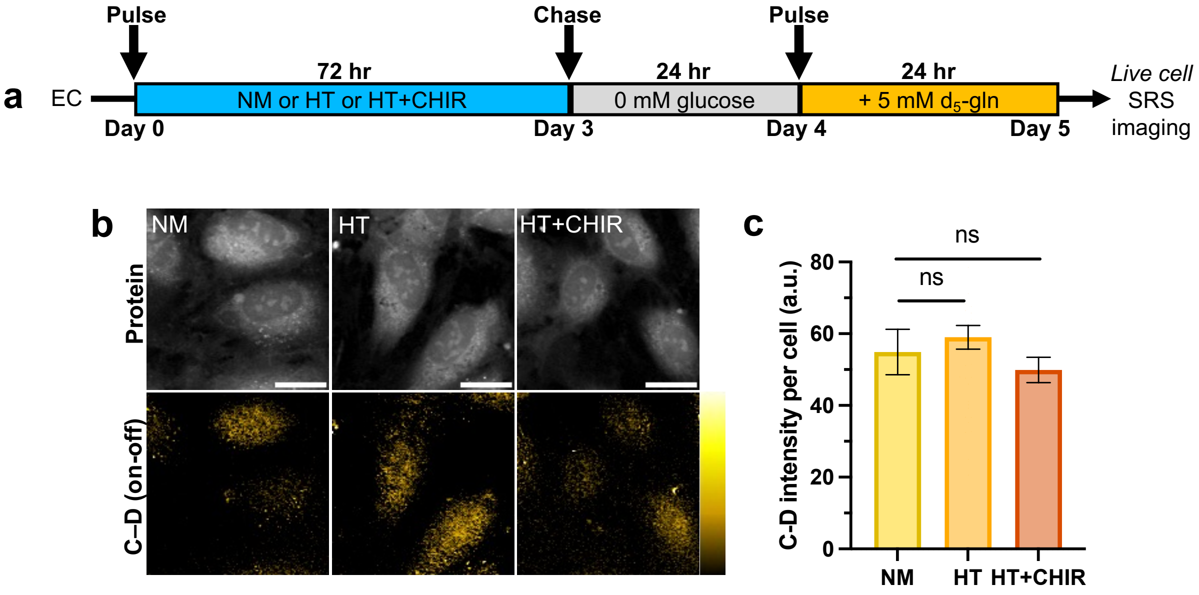


**Figure S11.** No differences in glutamine reliance upon glycogen depletion in ECs. **(a)** Illustration of pulse-chase experiment. After 72 hr of corresponding treatments, HUVECs were starved for 24 hr in glucose-free media followed by a chase with 5 mM $d_5$-gln in glucose-free media for 24 hr; **(b)** Representative SRS images at the Protein (–$CH_3$) and C–D (on-off) channels. Scale bar: 20 μm; **(c)** Quantitative analysis of SRS signal of C–D bonds per cell (n=38, 42, 39 cells for NM, HT, and HT+CHIR, respectively). Data are presented as mean ± SEM. Statistical significance was analyzed using two-tailed unpaired Student's t-tests, ns: no statistical difference.

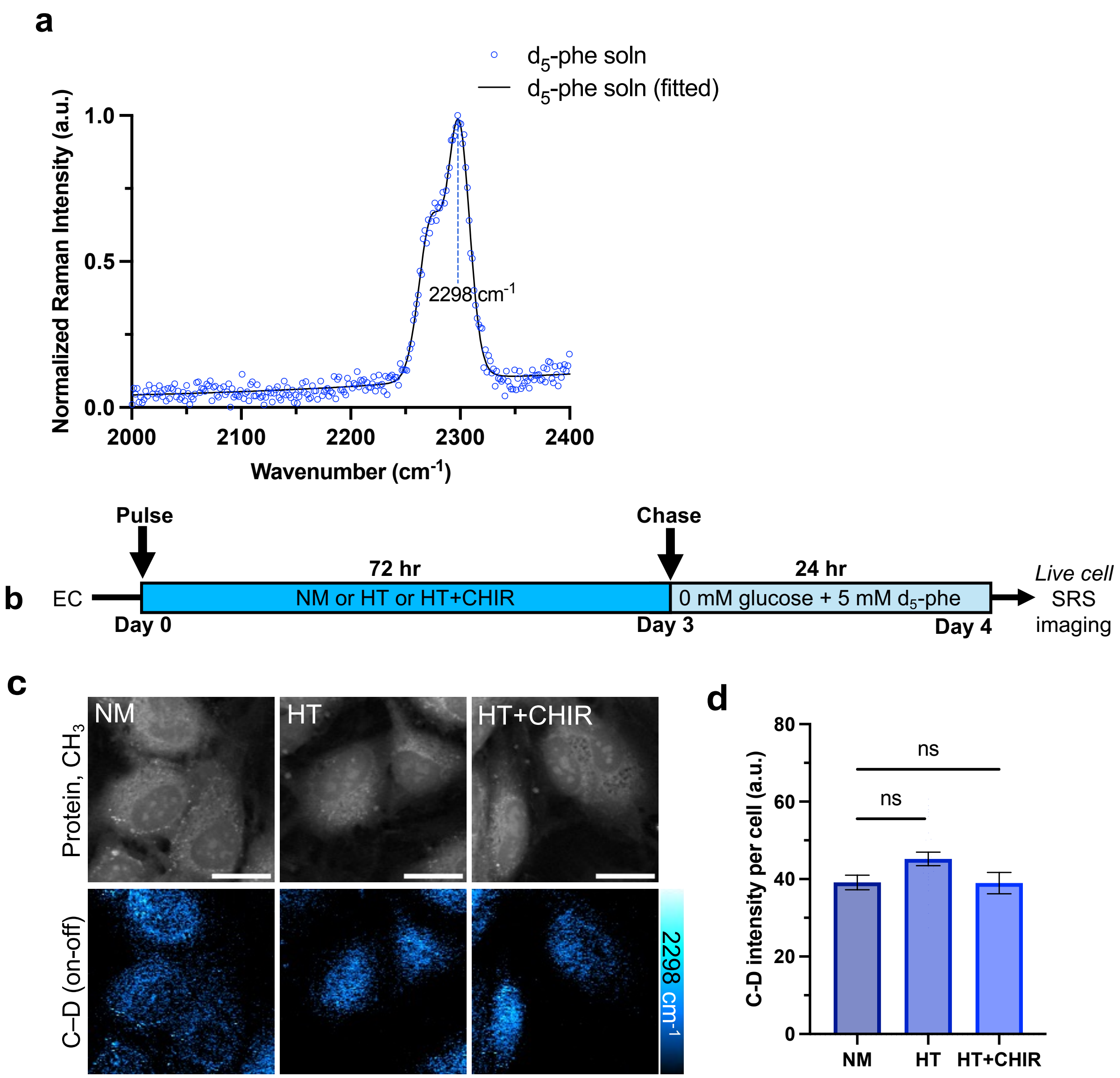


**Figure S12.** Presence of glycogen influences specific metabolic demands in ECs. **(a)** Normalized spontaneous Raman spectrum of 50 mM $d_5$-phenylalanine solution (soln) in water (experimental, blue circles; fitted, blue solid line); **(b)** Schematic of pulse-chase experiment with 5 mM $d_5$-phenylalanine in glucose-free ECGM. **(c)** Representative SRS images at the Protein (–$CH_3$) and C–D (on-off) channels. Scale bar: 20 μm; **(d)** Quantitative analysis of SRS signal of C–D bonds per cell (n=33, 25, 28 cells for NM, HT, and HT+CHIR, respectively). Data are presented as mean ± SEM. Statistical significance was analyzed using two-tailed unpaired Student's t-tests, ns: no statistical difference.

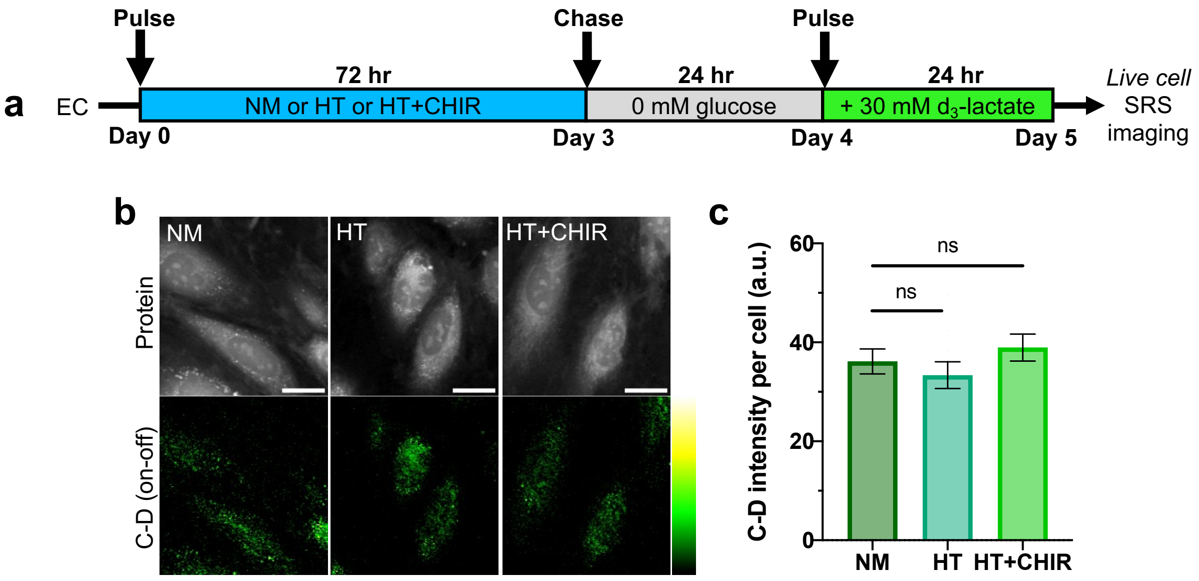


**Figure S13.** No differences in lactate reliance upon glycogen depletion. **(a)** Schematic of pulse-chase experiment. After 72 hr of corresponding treatments, HUVECs were starved for 24 hr in glucose-free media followed by a chase with 30 mM $d_3$-lac in glucose-free media for 24 hr; **(b)** Representative SRS images at the Protein ($–CH_3$) and C–D (on-off) channels. Scale bar: 20 μm; **(c)** Quantitative analysis of SRS signal of C–D bonds per cell (n=16, 14, 11 cells for NM, HT, and HT+CHIR, respectively). Data are presented as mean ± SEM. Statistical significance was analyzed using two-tailed unpaired Student's t-test, ns: no statistical difference.
